# Throughput and Resolution with a Next Generation Direct Electron Detector

**DOI:** 10.1101/620617

**Authors:** Joshua H. Mendez, Atousa Mehrani, Peter Randolph, Scott Stagg

## Abstract

Direct electron detectors (DEDs) have revolutionized cryo-electron microscopy by facilitating correction of beam-induced motion and radiation damage and by providing high-resolution image capture. A new generation of DEDs has been developed by Direct Electron, the DE64, that has good performance in both integrating and counting modes. Integrating mode is superior in throughput while counting mode is superior in image quality. We show that despite being ~10x slower in throughput, counting mode is superior in terms of reconstruction resolution per unit time of data collection.

## Main

The technology used for recording images for cryo-electron microscopy (cryo-EM) has evolved from the use of film to charge couple devices (CCD) and recently, direct electron detectors (DED) cameras. The progression of data collection techniques has allowed for the automation of data collection, increased signal to noise ratio for images, and for the correction of beam-induced motion. Direct detectors can operate in one of two modes: integrating mode, where electron hits deposit some amount of charge that is built up over the course of an exposure, and counting mode, where individual electron hits are “counted”, and the counts are summed over the exposure. Counting mode eliminates the Landau noise that results from individual electrons depositing different amounts of energy on the detector. On the other hand, counting requires a very weak beam and long exposure times which can result in lower throughput. These detector advancements together with the development of new software, have helped cryo-EM to break the 3 Å resolution boundary (Herzik, Wu, and Lander 2017). A new generation DED called the DE64 built by Direct Electron promises good performance when used in either integrating or counting mode. Resolution in single particle reconstructions is related to the amount of data contributing the reconstruction (Stagg et al. 2014; Heymann 2019), therefore the question arises, is it better to collect faster potentially lower-quality data in integrating mode or slower higher-quality data in counting mode.

Here we have characterized the DE64 and compared its performance when used in counting mode and in integrating mode. We have characterized its imaging performance by calculating the detective quantum efficiency (DQE) (Rose 1946) for both imaging modes and have used the DQE estimates to optimize the dose rate for counting. Imaging quality on actual samples was quantified by estimating Thon rings of carbon images and by reconstructing two different cryo-EM samples to better than 3 Å resolution. Finally, the imaging modes were compared in terms of resolution per unit time by comparing the resolution of sets of apoferritin particles collected in integrating mode, integrating mode with a Volta phase plate, and in counting mode.

The standard method to quantify the imaging power of a detector is to calculate its DQE. DQE is defined as 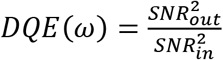 where ω is the spatial frequency. These are difficult quantities to measure experimentally, so DQE can be reformulated in terms of the camera’s modular transfer function (MTF) and its noise power spectrum (NPS), using the equation 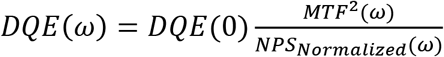 (McMullan et al. 2009). Thus to calculate the DE64’s DQE values, we first calculated its MTF using FindDQE (Ruskin, Yu, and Grigorieff 2013) and pair of images consisting in one flat field image with the beam blocker inserted half-way and the second image without the beam blocker inserted (Fig. 1A, Supplemental Fig. 1). When the MTFs for integrating and counting modes were compared, two different behaviors were observed. In integrating mode, the MTF decreased nearly linearly with increasing spatial frequency. The counting mode MTF, on the other hand, was close to the ideal MTF (Fig. 1A, purple). The NPS of for the different modes were calculated using an in-house python script. Images of the beam without sample were Fourier transformed, squared, and radially averaged to produce a one-dimension NPS curve. The zero frequency NPS was normalized to one as described in (McMullan et al. 2009) (Fig. 1B). Since counting detectors binarize individual electron hits, they are sensitive to the electron dose rate, and are susceptible to so-called coincidence loss (Li et al. 2013) where multiple electron hits are recorded as one due to their coincidence in position or time. Coincidence loss is reflected in the NPS as a suppression of the low frequency power (Fig. 1C). DE64 in counting mode operates at a frame rate of 141 frames/second, and NPS were calculated with dose rates of 2, 2.86, and 3.39 e^−^/p/s. There was a clear trend where 3.39 e^−^/p/s produced a substantial depression of the power at NPS(0). The depression was lessened with 2 and 2.86 e^−^/p/s. We did not try lower dose rates as that would have required unreasonably long exposures. The final value required for the DQE calculation, DQE(0), was calculated as described in (McMullan et al. 2014) where 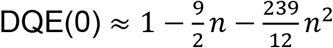 and *n* is the average electrons per frame per pixel. Given the MTF, NPS, and DQE(0), DQE curves were calculated for both integrating and counting modes demonstrating that they both show good performance over a broad range of frequencies. Both modes have similar values near Nyquist but differ at low frequencies where counting mode starts at a value of 0.93 and integrating starts at 0.52. These results were also reflected in the quality of Thon rings observed in images of carbon (Supplemental Fig. 2), where Thon rings were observed in both integrating and counting modes out to the Nyquist frequency, but there was better power in lower frequencies for counting mode. The better performance at lower frequencies for counting suggested that particles recorded in this mode can be better aligned and classified during single particle reconstruction.

**Figure 1:**
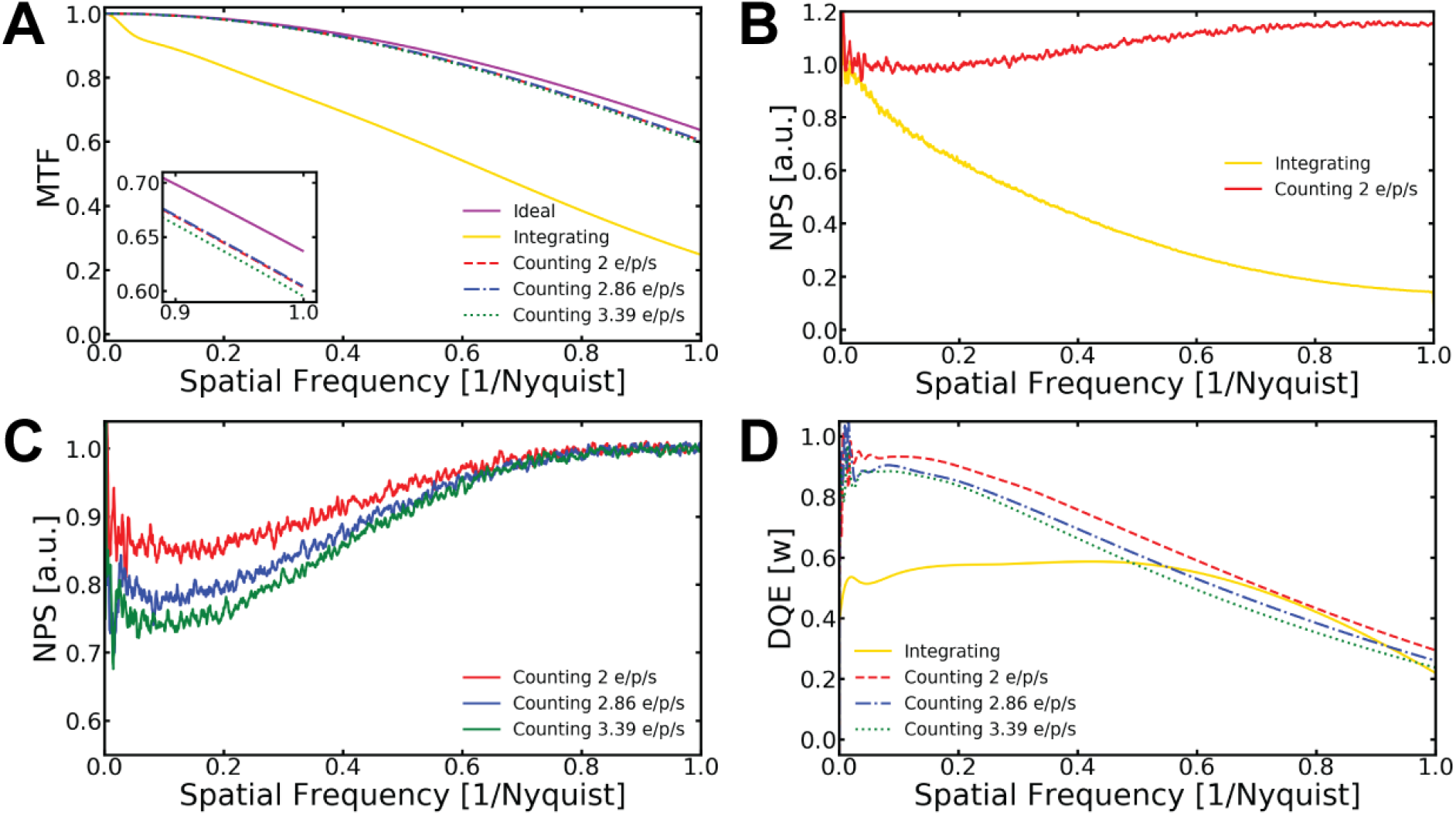
A) MTF curves of DE 64 in counting mode at different dose rates, integrating mode and the theoretical ideal curve. The inset is a zoom in view near Nyquist. B) DE 64 NPS curve comparison between integration and counting modes. C) NPS curves of DE 64 in counting mode at different dose rates. The NPS was normalized to a one for each dose rate. D) DQE curve of DE 64 in counting mode at different dose rates and integrating mode, where w is the spatial frequency.

**Figure 2:**
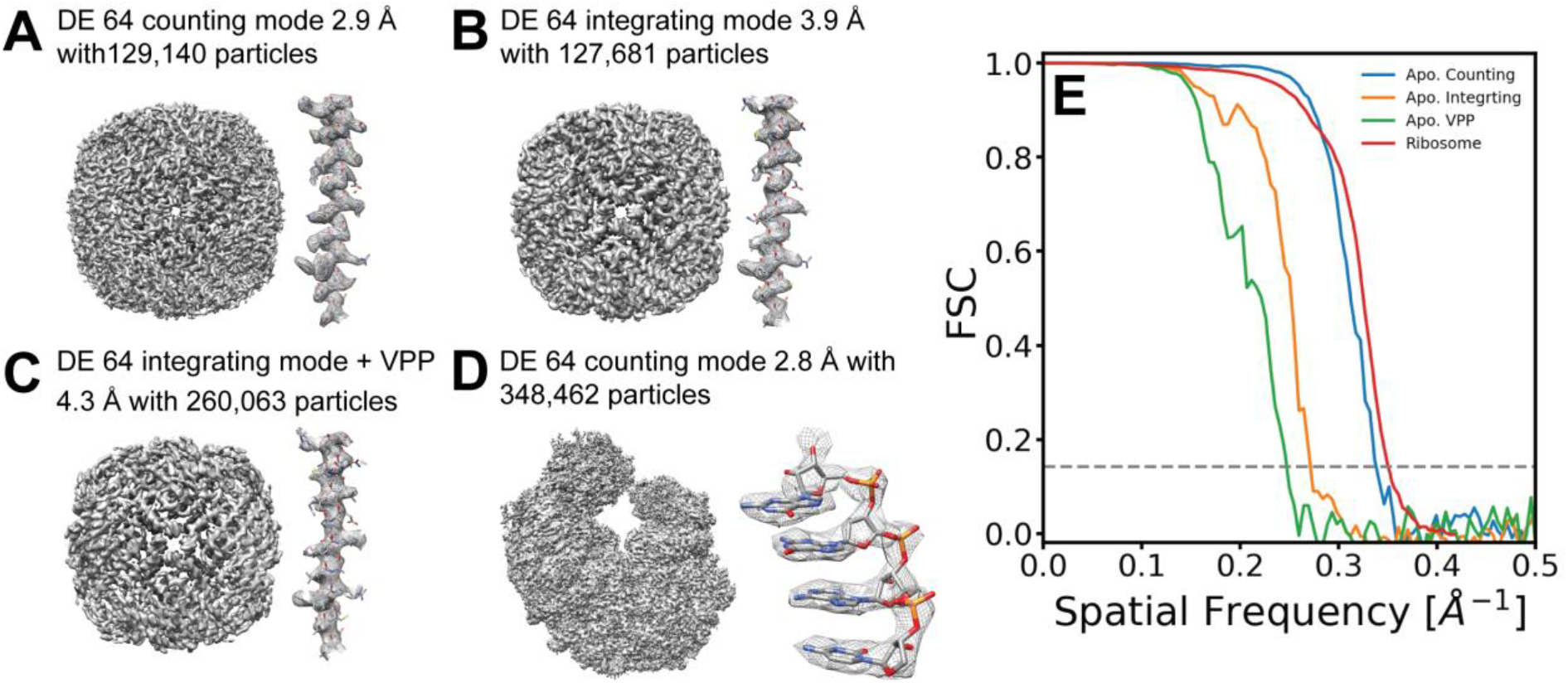
A) Counting mode density map with a resolution of 2.9 Å using 129,140 particles. B) Integration mode density map with a resolution of 3.9 Å using 127,681 particles. C) Integration mode using a Volta Phase-Plate density map with a resolution of 4.3 Å using 260,063 particles. D) Counting mode density map with a resolution of 2.8 Å using 348,462 particles. E) Fourier Shell Correlation of all reconstructions.

Since the resolution of cryo-EM reconstructions depends on both the quality and quantity of the images, the question arises, is it better to collect more data with integrating mode or better data with counting mode. We compared the camera modes by collecting and reconstructing data on the protein complex apoferritin as it can only be reconstructed with high quality images (Henderson and McMullan 2013; Russo and Passmore 2014; Massover 1993). In addition to the two camera modes, we also used integrating mode together with a Volta phase-plate (VPP) which boosts low frequency contrast. The idea here is that the combination of the boosted low frequency contrast from the VPP with the increased throughput for integrating mode might produce a better result than counting or integrating modes alone. We collected a total of 148,788 particles for counting, 152,220 particles for integrating, and 264,784 particles for integrating with VPP. These were refined using Relion resulting in reconstructions with resolutions of 2.9 Å for counting (EMD-20155), 3.9 Å for integrating (EMD-20156), and 4.3 Å for integrating with the VPP (EMD-20157) as seen in Figure 2. Clearly, despite the lower throughput, counting mode boosted the resolution that we were able to reach with approximately the same number of particles when compared to integrating mode. The increased low frequency contrast from the VPP did not help the resolution for integrating mode even with increased numbers of particles. Given that counting mode produced the best resolution on a per particle basis, we also characterized the performance of counting mode with an asymmetric particle. We collected 373,845 particles of a 70S ribosome sample. This was classified down to 348,462 particles and refined in Relion.

The resulting reconstruction achieved a resolution of 2.8 Å (EMD-20158) with well resolved features for the RNA and protein (Fig. 2D). Notably, this resolution was at ~4/5 of the Nyquist frequency indicating that resolution was beginning to be limited by sampling and higher resolution could likely be achieved by increasing the magnification during data collection.

The preceding data clearly indicate that data quality is critical for getting to high resolution, but since resolution is dependent upon the number of particles going into a reconstruction (LeBarron et al. 2008; Rosenthal and Henderson 2003), data collection throughput is also a critical parameter. It has been shown that to a first approximation there is a linear relationship between spatial frequency and the log of the number of particles contributing to a reconstruction, and these can be plotted to generate a ResLog plot (Stagg et al. 2014). The y-intercept of a ResLog plot is related to the alignability of a given dataset and the slope is related to the quality of the imaging. We performed a ResLog analysis for each of the three methods of data collection to understand how the reconstruction resolutions evolve as more particles were collected. This resulted in several observations about the quality of the data for the three different modes of data collection we used. We expected the VPP integrating particles to be more alignable than integrating alone given the boost in low frequency when the VPP is used. However, ResLog plots, seen in Figure 3A, showed that they both share essentially the same y-intercept indicating that particles of both datasets are equally alignable. The ResLog slope however was less for VPP integrating than it was for integrating alone. It is unclear why that may be, but we speculate that this is due to the additional challenges of estimating the phase shift as well as defocus when using the VPP. It is possible that inaccuracies in those estimates is what was limiting the per particle improvement in resolution for that mode. The counting mode particles had the highest y-intercept indicating that they had higher alignability than the other two modes. The ResLog slope for counting, on the other hand, was less than that of integrating mode. It should be noted, however, that the integrating curve would not intersect the counting curve until more than thirteen million particles have been collected.

**Figure 3:**
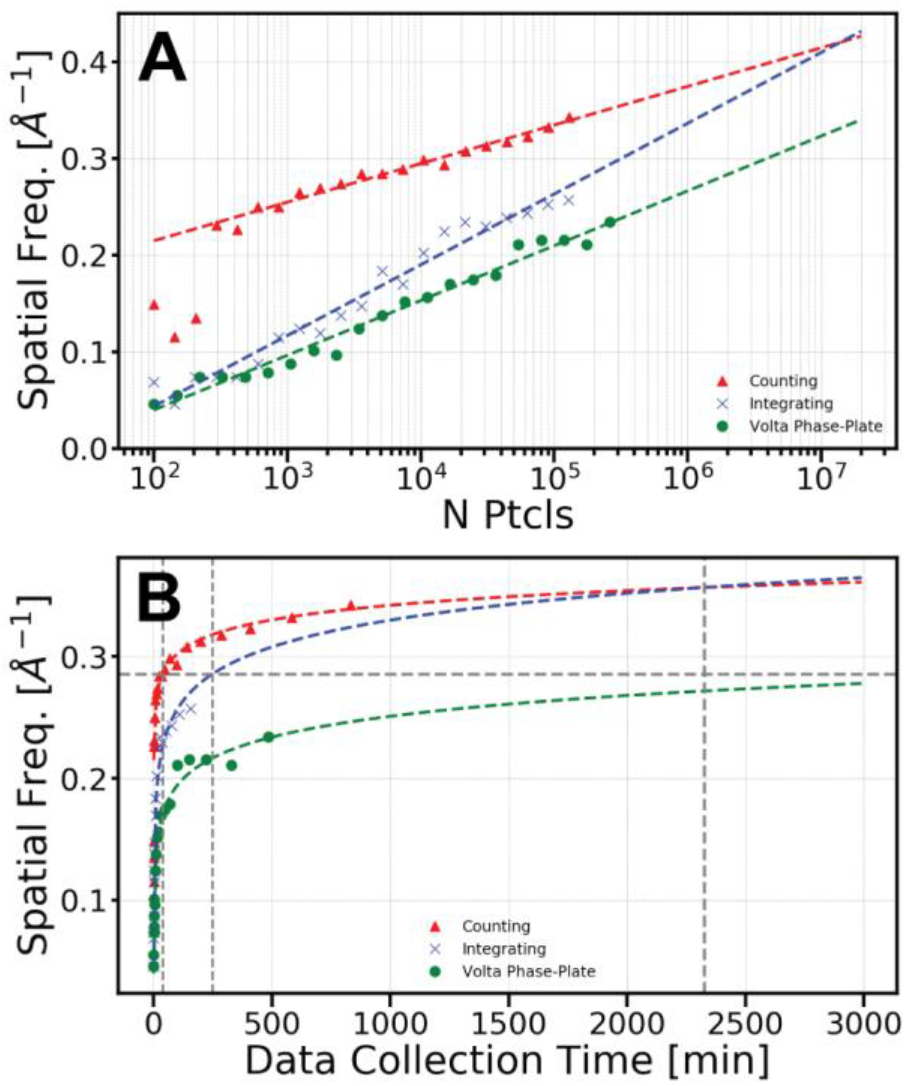
A) ResLog plots for apoferritin collected with counting, integrating, or integrating with VPP modes respectively. The integrating and counting modes intersect at 13 million particles. B) Plot of resolution against time of data collection. The horizontal line is the 3.5 Å resolution mark. The vertical lines are set at 38.7 minutes, 249.41 minutes and 2324.5 minutes.

The different modes of data collection for the DE64 have substantial differences in their data acquisition times. Integrating mode has the highest throughput, and the exposure time for a 61 e^−^/Å^2^ is 1.3 seconds. It also uses the full 8k × 8k sensor. In contrast, the exposure time for a 40 e^−^/Å^2^ in counting mode is 20.8 seconds, and the counting algorithm for the DE64 bins the image by two resulting in a 4k × 4k image. The acquisition time for a 30 e^−^/Å^2^ VPP integrating mode image is 4.6 seconds. Under real data collections with apoferritin, we obtained 42 micrographs per hour for integrating mode when using the one exposure per focus, resulting in 49,178 particles per hour. This can be increased to 113 micrographs or 132,312 particles per hour if we moved to four exposures per focus (Cheng et al. 2018) using image shift. In contrast, counting mode is limited to 36 micrographs or 9,349 particles per hour. The Volta-phase plate is limited to only a single exposure per focus since using image shift would result in an asymmetric buildup of charge on the VPP due to beam tilt. Acquisition with the VPP in integrating mode produced 27 micrographs or 32,203 particles per hour. When resolution was plotted against the time it took to acquire the particles for the different acquisition modes, counting mode was clearly superior in terms of resolution improvement over time. Remarkably, as seen in Figure 3B, it would take 4.2 hours to collect sufficient particles to achieve 3.5 Å resolution in integrating mode while that resolution could be achieved with particles collected in 38.7 minutes with counting. Thus, speed advantages of integrating mode are dwarfed by the increases in image quality with counting mode. Altogether, our data show that the DE64 shows potential to produce high resolution maps in both integrating and counting modes. The high throughput of integrating mode allows for the collection of large numbers of particles in a short period of time. Counting mode, although slower, achieved better resolutions whether comparing the same number of particles or the same amount of time. The large resolution improvements for counting compared to integrating suggest that its high DQE will enable structure determination for challenging single particle projects.

## Acknowledgements

We thank the New York Structural Biology Consortium for providing the apoferritin grid. We thank Dr. Christine Dunham and Dr. Eric Hoffer for providing the ribosome sample. Their work was supported by the NIH, R01GM093278. The work presented in this manuscript was supported by the NIH, R01GM108753. We also would like to extend our gratitude to Benjamin Bammes and everyone in Direct Electron for their assistance with DE64 counting mode.

## Methods

### Samples and Grid Preparation

The apoferritin grid was provided to us by New York Structural Biology Center (NYSBC). The sample was prepared by placing equine spleen apoferritin from Sigma-Aldrich onto a UltrAuFoil Quantifoil mesh grid. The grid was blotted, plunged, and stored for later use.

The second grid contained *E. coli* 70S ribosome cross-linked with KKL-2098 that was provided by the Dunham lab at Emory University. A 3 μl aliquot of ribosomes at a concentration of 100 nM was placed on plasma-cleaned UltrAuFoil® grids (Quantifoil, R1.2/1.3) and vitrified using Vitrobot Mark IV (FEI company). EM grids had been glow-discharged for 20 s using plasma cleaner (Gatan solarus 950). The Vitrobot chamber condition had been set to 100% humidity, 8°C temperature and 3.5 s blot time, and the cryogen used for vitrification was ethane cooled with liquid nitrogen.

### Data Collection

All data was collected using a DE 64 direct electron detector in conjunction with a FEI Titan Krios microscope set to a voltage of 300 kV and Leginon (Suloway et al. 2005). Apoferritin data was collected using counting mode, integrating mode and integrating mode with a Volta phase-plate (VPP). In counting mode, we used a magnification of 75000x with a calibrated pixel size of 0.928 Å. The total dose for each exposure was 40 e^−^/Å^2^ collected at a dose rate of 2.2 e^−^/Å^2^/s and a random defocus range from 0.5 to 3.0 μm. Raw frames were collected at rate of 141 frames/s and “quantized” (grouped) into movie frames containing an average of 1 e^−^/p. For integrating mode, we used a magnification of 37000x with a calibrated pixel size of 0.973 Å. Exposures were taken with a defocus ranging from 1.0 to 2.5 μm underfocus and a total electron dose of 61 e^−^/Å^2^. Frames were collected at rate of 32 frames/s with a dose rate of 50.6 e-/Å/s. Similarly, to integrating mode, we used a magnification of 37000x with a calibrated pixel size of 0.973 Å and a frame rate of 32 frames/s with VPP. In this case we used a smaller defocus ranged from 0.5 to 0.75 μm with an electron dose of 30 e^−^/Å^2^. With VPP we used a dose rate of 7 e^−^/Å^2^/s for optimal plate charging rate. The *E. coli* 70S Ribosome cross-linked with KKL-2098 was collected in counting mode at a dose rate of 1.3 e^−^/Å^2^/s. We used a magnification of 59000x with a calibrated pixel size of 1.19 e^−^/Å^2^. A total electron dose of 25 e^−^/Å^2^ and a defocus range from 1.5 to 3.0 μm was also used.

### Data Processing

During data acquisition micrographs were simultaneously processed using Appion (Lander et al. 2009), frame aligned, and CTF estimated. Movie frames were aligned using MotionCor2 (Zheng et al. 2017) and GCTF v1.06 (Zhang 2016) and CTFFIND4 (Rohou and Grigorieff 2015, 4) were used to estimate the CTF of the aligned micrographs. Particles were template picked from the aligned micrograph with FindEM (Roseman 2004). From the apoferritin sample using counting mode we collected 574 micrographs. Discriminating based on CTF confidence, 129,140 particles were extracted from micrographs with CTFs better than or equal to 4 Å resolution at the 0.5 cross-correlation coefficient mark. We collected 133 micrographs using integrating mode. Similarly, we extracted 127,681 particles from micrographs having CTFs better than or equal to 6.5 Å resolution at the 0.5 cross-correlation coefficient mark. When using integrating mode with VPP, we collected 222 micrographs extracting 260,063 particles. All particle where extracted using 224-pixel boxes, corresponding to approximately 1.5 times the size of the particles.

All refinements for apoferritin data were performed using Relion 3 with GPU acceleration. We did not do any classification to the particles. This includes, 2D classification or 3D classification. Our initial step was doing an initial reconstruction using all the particles. The initial model was created by downloading the final structure from EMPIAR-10026 and low-pass filtering it to a resolution of 20 Å. Other parameters used during the refinement process was the enforcement of octahedral symmetry and a mask diameter of 142 Å. After refinement, a custom mask was created using Relion; this was expanded by 5 voxels and a 7 voxel Gaussian decline was applied. The reconstructed map was sharpened using the post-processing function of Relion 3. Lastly, beam tilt estimation and correction was performed within Relion 3 CTF refinement (Zivanov et al. 2018). We continued the series of correction until no improvement was seen. The reconstructed maps of apoferritin in counting mode, integrating mode, and integrating mode with VPP were deposited in EMDB with the IDs EMD-20155, EMD-20156, and EMD-20157, respectively.

For the ribosome sample, we collected 1,752 micrographs. Particles were extracted from all micrographs with a box size of 384 pixels creating a stack of 373,845 particles. The particles were initially processed with cisTEM (Grant, Rohou, and Grigorieff 2018). A 2D classification of the particles was performed only selecting classes showing defined features. The classified particles were refined and further classified using 3D classification. A final reconstruction was performed with Relion 3. Within Relion 3 beam-tilt estimations and corrections were performed until no improvement was seen. The map was deposited in EMDB with the ID, EMD-20158.

## References

Cheng, Anchi, Edward T. Eng, Lambertus Alink, William J. Rice, Kelsey D. Jordan, Laura Y. Kim, Clinton S. Potter, and Bridget Carraggher. 2018. “High Resolution Single Particle Cryo-Electron Microscopy Using Beam-Image Shift.” BioRxiv, April, 306241. https://doi.org/10.1101/306241.

Henderson, Richard, and Greg McMullan. 2013. “Problems in Obtaining Perfect Images by Single-Particle Electron Cryomicroscopy of Biological Structures in Amorphous Ice.” Microscopy 62 (1): 43–50. https://doi.org/10.1093/jmicro/dfs094.

Herzik, Mark A., Mengyu Wu, and Gabriel C. Lander. 2017. “Achieving Better than 3 Å Resolution by Single Particle Cryo-EM at 200 KeV.” Nature Methods 14 (11): 1075–78. https://doi.org/10.1038/nmeth.4461.

Heymann, J. B. 2019. “Single-Particle Reconstruction Statistics: A Diagnostic Tool in Solving Biomolecular Structures by Cryo-EM.” Acta Crystallographica Section F: Structural Biology Communications 75 (1): 33–44. https://doi.org/10.1107/S2053230X18017636.

LeBarron, Jamie, Robert A. Grassucci, Tanvir R. Shaikh, William T. Baxter, Jayati Sengupta, and Joachim Frank. 2008. “Exploration of Parameters in Cryo-EM Leading to an Improved Density Map of the E. Coli Ribosome.” Journal of Structural Biology 164 (1): 24–32. https://doi.org/10.1016/j.jsb.2008.05.007.

Li, Xueming, Shawn Q. Zheng, Kiyoshi Egami, David A. Agard, and Yifan Cheng. 2013. “Influence of Electron Dose Rate on Electron Counting Images Recorded with the K2 Camera.” Journal of Structural Biology 184 (2): 251–60. https://doi.org/10.1016/j.jsb.2013.08.005.

Massover, William H. 1993. “Ultrastructure of Ferritin and Apoferritin: A Review.” Micron 24 (4): 389–437. https://doi.org/10.1016/0968-4328(93)90005-L.

McMullan, G., S. Chen, R. Henderson, and A. R. Faruqi. 2009. “Detective Quantum Efficiency of Electron Area Detectors in Electron Microscopy.” Ultramicroscopy 109 (9): 1126–43. https://doi.org/10.1016/j.ultramic.2009.04.002.

McMullan, G., A. R. Faruqi, D. Clare, and R. Henderson. 2014. “Comparison of Optimal Performance at 300keV of Three Direct Electron Detectors for Use in Low Dose Electron Microscopy.” Ultramicroscopy 147 (December): 156–63. https://doi.org/10.1016/j.ultramic.2014.08.002.

Rose, A. 1946. “A Unified Approach to the Performance of Photographic Film, Television Pickup Tubes, and the Human Eye.” Journal of the Society of Motion Picture Engineers 47 (4): 273–94. https://doi.org/10.5594/J12772.

Rosenthal, Peter B., and Richard Henderson. 2003. “Optimal Determination of Particle Orientation, Absolute Hand, and Contrast Loss in Single-Particle Electron Cryomicroscopy.” Journal of Molecular Biology 333 (4): 721–45. https://doi.org/10.1016/j.jmb.2003.07.013.

Ruskin, Rachel S., Zhiheng Yu, and Nikolaus Grigorieff. 2013. “Quantitative Characterization of Electron Detectors for Transmission Electron Microscopy.” Journal of Structural Biology 184 (3): 385–93. https://doi.org/10.1016/j.jsb.2013.10.016.

Russo, Christopher J., and Lori A. Passmore. 2014. “Ultrastable Gold Substrates for Electron Cryomicroscopy.” Science 346 (6215): 1377–80. https://doi.org/10.1126/science.1259530.

Stagg, Scott M., Alex J. Noble, Michael Spilman, and Michael S. Chapman. 2014. “ResLog Plots as an Empirical Metric of the Quality of Cryo-EM Reconstructions.” Journal of Structural Biology 185 (3): 418–26. https://doi.org/10.1016/j.jsb.2013.12.010.

## Methods References

Grant, Timothy, Alexis Rohou, and Nikolaus Grigorieff. 2018. “CisTEM, User-Friendly Software for Single-Particle Image Processing.” Edited by Edward H Egelman. ELife 7 (March): e35383. https://doi.org/10.7554/eLife.35383.

Lander, Gabriel C., Scott M. Stagg, Neil R. Voss, Anchi Cheng, Denis Fellmann, James Pulokas, Craig Yoshioka, et al. 2009. “Appion: An Integrated, Database-Driven Pipeline to Facilitate EM Image Processing.” Journal of Structural Biology 166 (1): 95–102. https://doi.org/10.1016/j.jsb.2009.01.002.

Rohou, Alexis, and Nikolaus Grigorieff. 2015. “CTFFIND4: Fast and Accurate Defocus Estimation from Electron Micrographs.” Journal of Structural Biology, Recent Advances in Detector Technologies and Applications for Molecular TEM, 192 (2): 216–21. https://doi.org/10.1016/j.jsb.2015.08.008.

Roseman, A. M. 2004. “FindEM—a Fast, Efficient Program for Automatic Selection of Particles from Electron Micrographs.” Journal of Structural Biology, Automated Particle Selection for Cryo-Electron Microscopy, 145 (1): 91–99. https://doi.org/10.1016/j.jsb.2003.11.007.

Suloway, Christian, James Pulokas, Denis Fellmann, Anchi Cheng, Francisco Guerra, Joel Quispe, Scott Stagg, Clinton S. Potter, and Bridget Carragher. 2005. “Automated Molecular Microscopy: The New Leginon System.” Journal of Structural Biology 151 (1): 41–60. https://doi.org/10.1016/j.jsb.2005.03.010.

Zhang, Kai. 2016. “Gctf: Real-Time CTF Determination and Correction.” Journal of Structural Biology 193 (1): 1–12. https://doi.org/10.1016/j.jsb.2015.11.003.

Zheng, Shawn Q., Eugene Palovcak, Jean-Paul Armache, Kliment A. Verba, Yifan Cheng, and David A. Agard. 2017. “MotionCor2: Anisotropic Correction of Beam-Induced Motion for Improved Cryo-Electron Microscopy.” Nature Methods 14 (4): 331–32. https://doi.org/10.1038/nmeth.4193.

Zivanov, Jasenko, Takanori Nakane, Björn O Forsberg, Dari Kimanius, Wim JH Hagen, Erik Lindahl, and Sjors HW Scheres. 2018. “New Tools for Automated High-Resolution Cryo-EM Structure Determination in RELION-3.” Edited by Edward H Egelman and John Kuriyan. ELife 7 (November): e42166. https://doi.org/10.7554/eLife.42166.

