## Supplemental Material for "Throughput and Resolution with a Next Generation Direct Electron Detector"

### Supplementary

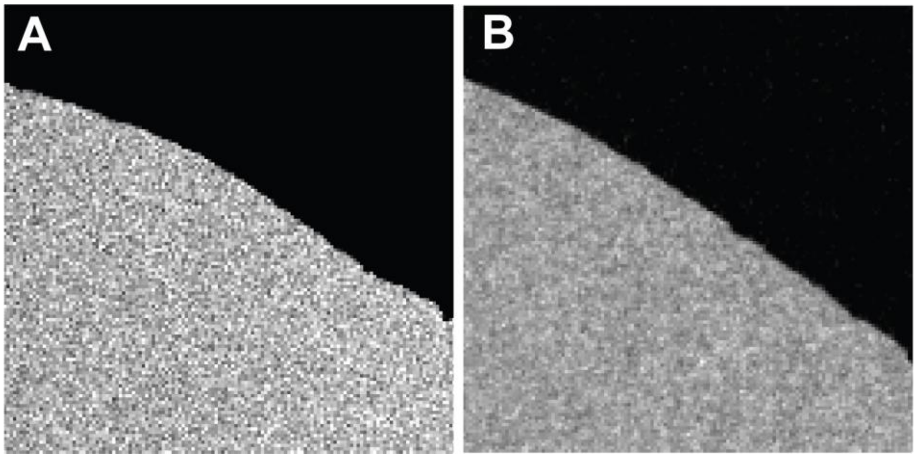


**Figure 1:** A) Beam stop sharp edge imaged in counting mode. B) Beam stop sharp edge imaged in integrating mode

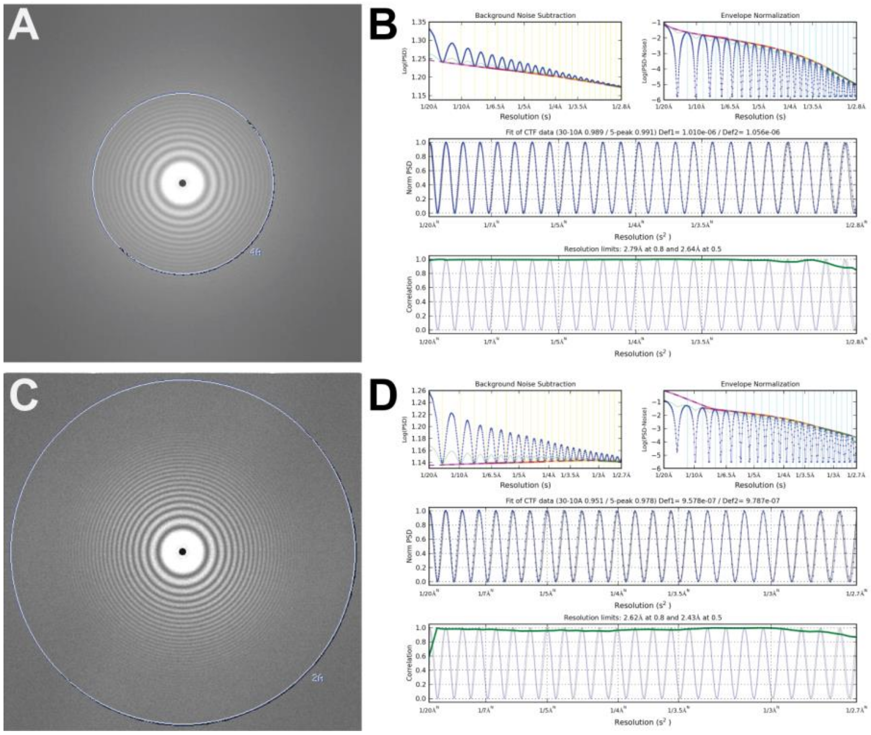


**Figure 2:** A) Thon rings of carbon from an image acquired using DE 64 in integration mode. B) 1-dimension radial average of the power spectrum with the fitted contrast transfer function (CTF) as seen in Appion. C) Thon rings of carbon from an image acquired using DE 64 in counting mode. D) 1-dimension radial average of the power spectrum with the fitted CTF as seen in Appion.
